# Genomic prediction including SNP-specific variance predictors

**DOI:** 10.1101/636746

**Authors:** E. F. Mouresan, M. Selle, L. Rönnegård

**Affiliations:** Department of Animal Breeding and Genetics, Swedish University of Agricultural Sciences, Sweden, 75007; Department of Mathematical Sciences, Norwegian University of Science and Technology, Norway, 7491; School of Technology and Business Studies, Dalarna University, Sweden, 79188

**Keywords:** Genomic selection, BLUP, hglm, CodataGS, external information

## Abstract

The amount of available biological information on the markers is constantly increasing and provides valuable insight into the underlying biology of traits of interest. This information can also be used to inform the models applied for genomic selection to improve predictions. The objective of this study was to propose a general model for genomic selection using a link function approach within the hierarchical generalized linear model framework (hglm) that can include external information on the markers. These models can be fitted using the well-established hglm package in R. Furthermore, we also present an R package (CodataGS) to fit these models, which is significantly faster than the hglm package when the number of markers largely exceeds the number of individuals. Simulated data was used to validate the proposed model. Knowledge on the location of the QTLs on the genome, with varying degree of uncertainty, was used as external information on the markers. The proposed model showed improved accuracies from 3.8% up to 23.2% compared to the SNP-BLUP method, which assumes equal variances for all markers. The performance of the proposed models depended on the genetic architecture of the trait, as traits that deviate from the infinitesimal model benefited more from the external information. Also, the gain in accuracy depended on the degree of uncertainty of the external information provided to the model. The usefulness of these type of models is expected to increase with time as more accurate information on the markers becomes available.

## Introduction

The identification of a large number of Single Nucleotide Polymorphisms (SNPs) along the genome, as a by-product of the sequencing efforts (Daetwyler et al., 2014) and the development of SNP-chip genotyping technology (Gunderson et al., 2005) have made genotyping of thousands of markers affordable at low cost. Meuwissen et al. (2001) could foresee these breakthroughs in technology and proposed a new method of selection in animal breeding denoted as Genomic Selection (GS). This method has been tested through simulation studies (Meuwissen et al., 2001; Muir, 2007) and cross validation with real data in different species such as mice (Legarra et al., 2008), dairy cattle (Luan et al., 2009; VanRaden et al., 2009), aquaculture (Sonesson and Meuwissen, 2009) and poultry (Gonzalez-Recio et al., 2009) and nowadays, GS has become part of the routine breeding schemes in dairy cattle (Hayes et al., 2009) and other species including pigs (Ostersen et al., 2011; Hidalgo et al., 2015; Tusell et al., 2016) and poultry (Wolc et al., 2015).

Several statistical models have been proposed for genomic prediction using whole-genome markers. The most popular method provides best linear unbiased prediction (BLUP) estimates of the SNP markers (Meuwissen et al., 2001) by assuming that the marker effects come from a Gaussian distribution with constant variance and results in all SNPs having some effect on the analysed trait. This method is referred to either as GBLUP or SNP-BLUP depending on the implementation, but independent of implementation the method still assumes that all markers have some effect. Biologically, it seems more reasonable to assume that some of the markers are in linkage disequilibrium (LD) with a causative gene or a quantitative trait locus (QTL) and therefore can capture their effect on the studied trait, whereas some markers are not in LD with any gene and should therefore not capture any effect. To achieve this idea, several methods have been developed to incorporate different prior assumptions on the genetic architecture of the trait. For this family of methods, often referred to as the Bayesian Alphabet (Gianola, 2013), it is assumed that the genetic effects of the SNPs follow alternative distributions like a t-distribution (Bayes A) (Meuwissen et al., 2001), a double exponential distribution (Bayes LASSO) (de los Campos et al., 2009; Usai et al., 2009) or a mixture of distributions (i.e. Bayes B, Bayes Cπ, Bayes R) (Meuwissen et al., 2001; Habier et al., 2011; Erbe et al., 2012). The prior assumptions of these methods are rather arbitrary and their performance relies heavily on the model assumptions capturing accurately the true genetic architecture of the trait of interest (Daetwyler et al., 2010; Hayes et al., 2010).

Whole-genome sequencing of individuals has facilitated the detection of genetic variants that can be used for GS. Currently, in *Bos Taurus* cattle ^~^28 million genetic variants have been reported (Daetwyler et al., 2013). This large number of polymorphic markers comes with a major challenge in terms of computational speed and memory. One way to deal with this problem is to make use of the biological information available on the markers, e.g. to annotate the markers in classes based on genome location or functionality and prioritize those classes that show a higher probability of containing trait associated markers. Koufariotis et al. (2014) showed that protein coding regions explain significantly more variation than similar number of random chosen markers across many traits in cattle. Moreover, the upstream and downstream classes also showed significant enrichment in trait associated variants suggesting that these classes can potentially have important regulatory functions. In the same line, Yang et al. (2011) stated that genic regions contributed more additive genetic variance than non-genic regions for human traits. However, Do et al. (2015) found that the contribution to total genomic variance per SNP among the annotated classes was similar for all regions in a feed efficiency study in pigs.

Several authors have also investigated the predictive ability of models based on annotation classes. Using kernel methods, Morota et al. (2014) and Abdollahi-Arpanahi et al. (2016) showed that a whole-genome approach provided better predictive ability than that obtained from classes of genomic regions considered separately. Likewise, Do et al. (2015) using GBLUP and Bayesian methods (Bayes A, B and Cπ) found that classification of SNPs by genomic annotation had little impact on the accuracy of prediction for feed efficiency traits in pigs.

Apart from genome annotation information, other biological information is available on the SNPs. QTL databases are available for most livestock species (Hu et al., 2013) and Genome-Wide Association Studies (GWAS) (Bush and Moore, 2007) have identified a great number of trait-associated markers. Moreover, metabolic and signalling pathways (Kanehisa et al., 2008; Croft et al., 2011; Caspi et al., 2012) and gene regulatory networks (Lee et al., 2002; Shalgi et al., 2007; Hecker et al., 2009) can also provide valuable insight to the underlying biology of the traits of interest (Snelling et al., 2013). A rather new tool that has been developed to incorporate existing knowledge of the genetic architecture of complex traits into a GS model is BLUP|GA, i.e. “BLUP approach given the Genetic Architecture” (Zhang et al., 2014). This tool uses publicly available GWAS results and showed improved prediction accuracies compared to traditional GBLUP and Bayes B methods. Also, a similar approach was developed by Kadarmideen (2014) (system genomic BLUP, – sgBLUP-) where SNPs with known biological role were explicitly modelled in addition to conventional random SNP effects in SNP-BLUP or GBLUP methods. Along with the BLUP approaches, several Bayesian methods were also developed. Bayes Bπ (Gao et al., 2015) is a modified version of Bayes B (Meuwissen et al., 2001) able to utilize locus-specific priors. In their study, the authors obtained locus-specific priors from variance analysis (ANOVA) based on information from each single marker separately and the results showed improved accuracy and decreased bias compared to Bayes B and Bayes Cπ. In a similar way, MacLeod et al. (2016) proposed a modification to the BayesR method (Erbe et al., 2012) that incorporates prior biological knowledge. This method provides a flexible approach to improve the accuracy of genomic prediction and QTL discovery taking advantage of available biological knowledge. The basic idea of previously developed methods is to group SNPs into those having a biological function and those with an unknown function. Both the BLUP|GA and BayesBπ methods, also include continuous weights for all, or a subset of markers. For the BLUP|GA method, weights computed using trait-specific GWAS results are used to construct the genomic relationship matrix, whereas in BayesBπ the weights are computed from single-SNP ANOVA analyses.

Although a large number of methods have been developed already for GS, a general BLUP method to include explanatory variables for SNP-specific variances that allow both continuous and class variables seems to be missing. Here we propose a general model using a link function approach within the hierarchical generalized linear model framework (Lee et al., 2006). The algorithm proposed by Lee and Nelder (1996) is used, where the hierarchical generalized linear model is fitted by iterating between augmented generalized linear models. With this approach, rather complex models can be fitted using a single deterministic fitting algorithm (see Rönnegård et al. 2010a, 2010c).

The aim of the paper is to assess the accuracy for such a model including predictors for SNP variances, with special emphasis on the effect of the trait’s genetic architecture and LD structure on estimation accuracy. We present a family of models where the SNP variances can be modelled using both, categorical and continuous predictors, or a combination of the two. The computation time of these models is also studied and a new, faster R package (CodataGS) to fit these models is presented.

## Materials and Methods

### Data simulation

Data was simulated to evaluate the models. Four different scenarios for QTL variance distribution were simulated under three different genetic architectures in which the number of QTL per chromosome was 10, 20 or 100. For each combination of scenario and genetic architecture, 100 simulation replicates were produced. This section describes the simulations in detail.

A base population was simulated of 100 individuals that evolved under random mating for 400 non-overlapping generations (generation −399 to 0) maintaining the population size constant. After the 400 historical generations, two more generations were simulated, still under random mating and expanding the population size from 100 to 200 individuals per generation. Generation 1 was used as training set and generation 2 as validation set. The genome comprised of two chromosomes of 1 Morgan each with 8,800 loci, evenly distributed across the genome. In the base population alleles were coded as 0 or 1 with equal probability resulting in intermediate average allele frequencies. In the first generation, 1,000 loci per chromosome were selected randomly among those loci with a Minor allele frequency (MAF) higher than 0.05 to simulate the SNP marker panel. The same loci were used for validation in generation 2.

To simulate phenotypes in generation 1 (training set), *N_QTL_* loci were selected randomly excluding loci that were on the edge of the chromosome and those with a MAF lower than 0.05. In order to simulate different scenarios of genetic architecture underlying the trait, the number of QTLs (*N_QTL_*) varied between 10, 20 and 100 per chromosome. Moreover, the QTL effects, *u_j_*, *j* = 1, …, *N_QTL_*, were assumed to be normally distributed with mean 0 and varying variance assigned in one of the following ways:

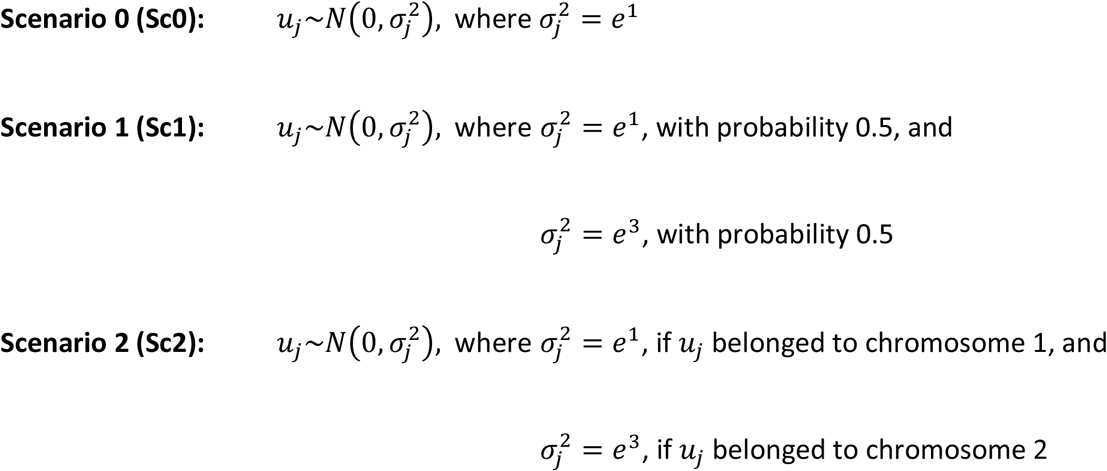

The difference between the scenarios Sc1 and Sc2 is that in Sc1 heterogeneous QTL effects are allowed on the same chromosome and may be in linkage disequilibrium with each other. On the other hand, in Sc2 the two different types of QTLs are located on different chromosomes to ensure low LD between them.

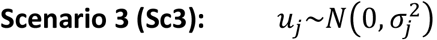, where 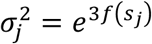, *s_j_* is the position of QTL *j* and *f* is a function of relative distance to the chromosome edge. Consequently, 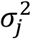 take values between *e*^1^ and *e*^3^.

For each scenario, the three separate genetic architectures were simulated, i.e. with 10, 20 or 100 QTL per chromosome. In order for the results from the different scenarios and genetic architectures to be comparable, the total genetic variance was scaled to 1.0. In this way, the obtained traits were either controlled by a small number of QTLs with medium-large effects or by a large number of QTLs with small effects.

In generation 1 (training set) phenotypes were simulated for all 200 individuals as:

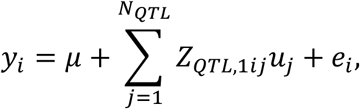

where *y_i_* is the phenotype of individual *i*, *μ* is a fixed effect which was set equal to 0, *Z*_*QTL*,1*ij*_ is the genotype for the *j^th^* QTL coded as 0, 1 or 2 for the homozygote, heterozygote and the alternative homozygote respectively for individual *i* in generation 1, *u_j_* is the simulated normally distributed *j^th^* QTL effect as described above, and *e_i_* is the residual effect of the *i^th^* individual normally distributed with mean 0 and the appropriate variance 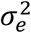 in order to create a trait with heritability of 0.2.

Generation 2 was used as validation set where true genomic breeding values (TGBVs) were computed as:

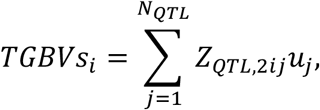

where *Z*_*QTL*,2*ij*_ is the QTL genotype for QTL *j* and individual *i* for this generation.

### Genomic evaluation

To estimate the SNP effects, the marker panel of 1,000 SNPs per chromosome mentioned above was used and the following model was assumed:

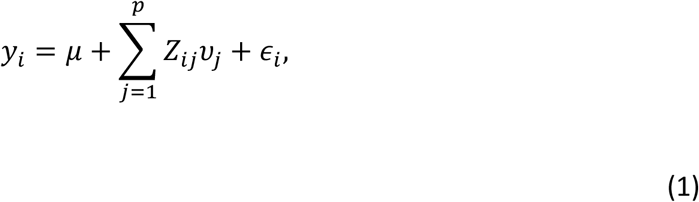

where *y_i_* is the phenotype of individual *i*, *μ* is a fixed effect, *p* is the total number of SNPs, *Z_ij_* is the genotype of the SNP *j* for individual *i* coded as 0, 1 or 2, 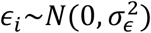 is the residual effect, and

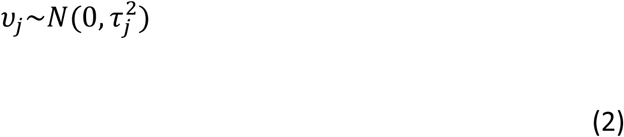

is the *j^th^* SNP effect normally distributed with mean 0 and variance

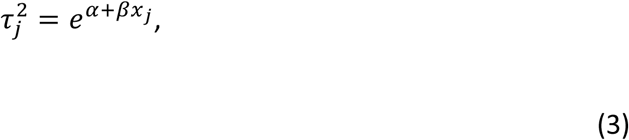

where *α* + *βx_j_* is a linear predictor for the SNP-specific variances.

### Evaluation models

The linear predictor (*α* + *βx_j_*) allows to incorporate any type of external information about the SNP variance, making it possible to assign the same variance for all SNPs, a subgroup of SNPs or assign a unique variance for each SNP. We used this linear predictor to introduce external information on the SNPs into the models and the predictive performance of different prior assumptions was tested. The models tested in this study were:

1. **SNP-BLUP**: In the traditional model the variances of the markers are assumed to be equal for all markers and therefore *x_j_* = 1 in the linear predictor for all markers.
2. Categorical models (**W10, W20** and **W40**): For these models the genome was divided into non-overlapping windows of 10, 20 or 40 SNPs. Then, all the SNPs within a given window were given the value *x_j_* = 1 if they contained a QTL and *x_j_* = 0 if they did not. Hence, a study with known regions harbouring the QTL was mimicked, where these regions were known with varying degree of uncertainty.
3. Continuous model (**LD**): For this model, the linkage disequilibrium (LD) between a SNP and a QTL was calculated as *r*^2^ = *D*^2^/(*p_S_p_s_p_Q_p_q_*), where *D* = *f_SQ_f_sq_* – *f_Sq_f_sQ_* (Falconer and Mackay, 1996), *p_S_*, *p_s_*, *p_Q_* and *p_q_* are the allele frequencies of the SNP and QTL, *f_SQ_*, *f_sq_* are the homozygous haplotype frequencies and *f_Sq_*, *f_sQ_* are the heterozygous haplotype frequencies. Then, each SNP was assigned the value of 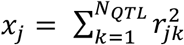. The relationship between SNPs and QTLs was modelled in such way that markers in higher LD with one or more QTLs would be given more importance in the model compared to other markers not in LD with any QTL.
4. Combination of categorical and continuous models (**W10-LD**, **W20-LD** and **W40-LD**): In these models the genome was divided into windows as in the previous categorical models but the SNPs located within a window that harboured a QTL were given the value of the LD with the QTL instead of 1. The model could, therefore, differentiate between SNPs not only based on location but also based on the relationship with the real QTL. Table 1 gives an overview of all simulated scenarios and models tested. Each scenario was simulated with 10, 20 and 100 QTL per chromosome as described previously.
5. Additional models (**W10-2var, W20-2var, W40-2var, Dis, W10-Dis, W20-Dis** and **W40-Dis**): The previously described models include external information on the physical location of the QTLs relative to the SNPs or/and the relationship of the SNPs with the QTLs but they do not include any information about the QTL variances. Therefore, a few additional models were created based on the particular parameters used for the simulation of each genetic architecture scenario. These models are defined as follows.

a. For the scenarios where the QTL effects came from distributions with two different variances (Sc1 and Sc2) we assumed this information was known and we expanded the linear predictor to *α* + *βx*_*j*1_ + *γx*_*j*2_ in order to accommodate for more variances (in the models W10-2var, W20-2var, and W40-2var). The genome was divided in non-overlapping windows as before and SNPs associated with a QTL with variance 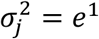 was assigned *x*_*j*1_ = 1 and *x*_*j*2_ = 0, while if it was associated with a QTL with variance 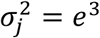 it was assigned *x*_*j*1_ = 0 and *x*_*j*2_ = 1. If a SNP was located within a window with no QTL then both *x*_*j*1_ and *x*_*j*2_ had a value of 0.
b. For Sc3, we used the distance of the markers from the edge of the chromosome as external information either as a continuous variable (Dis) or within windows (W10-Dis, W20-Dis and W40-Dis), since the QTL variances were simulated in the same way.

**Table 1.**
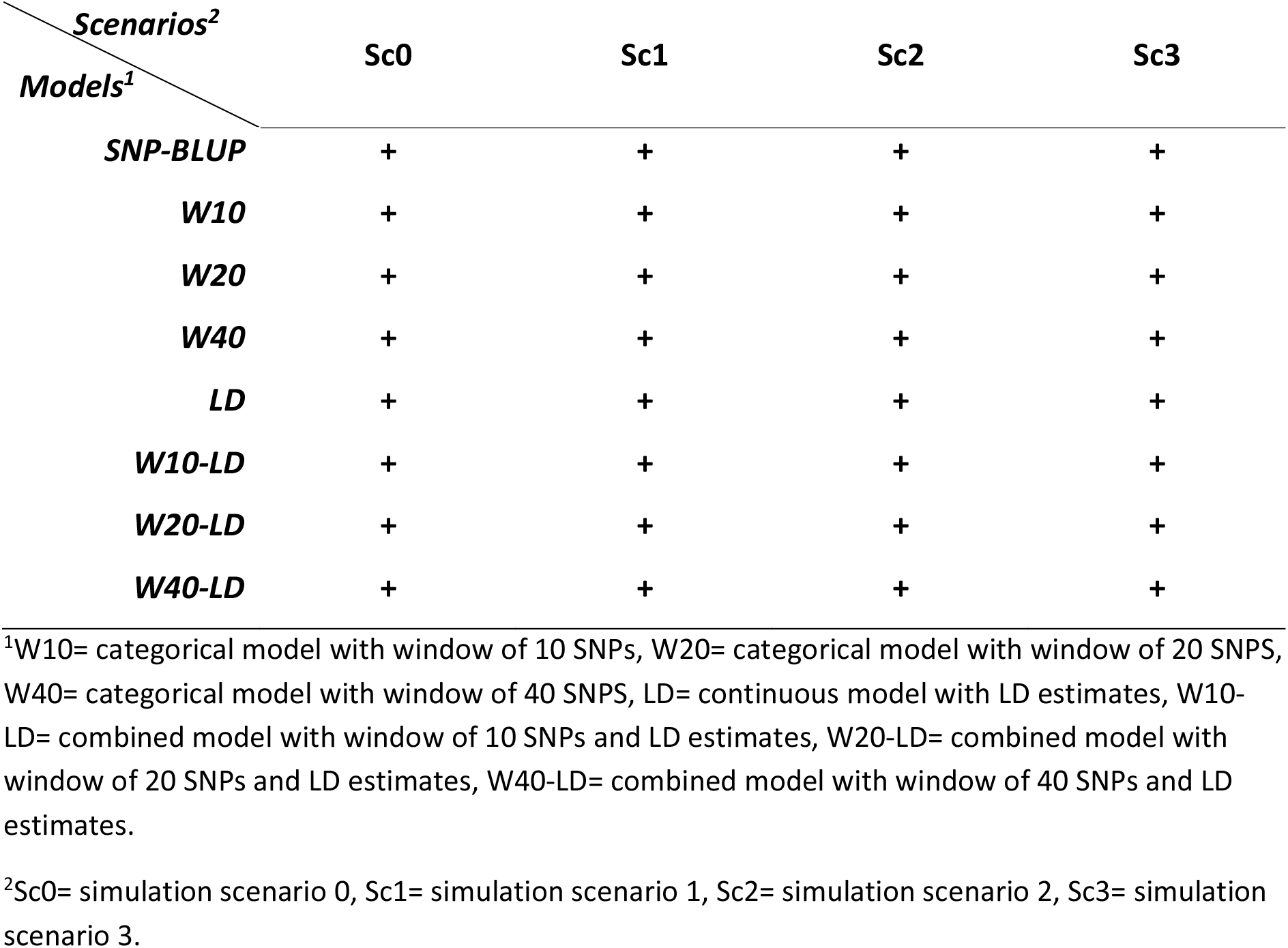
Summary of models tested for each scenario of genetic architecture simulated.

| *Models<sup>1</sup>* | *Scenarios<sup>2</sup>* |  |  |  |
| --- | --- | --- | --- | --- |
|  | Sc0 | Sc1 | Sc2 | Sc3 |
| ***SNP-BLUP*** | + | + | + | + |
| ***W10*** | + | + | + | + |
| ***W20*** | + | + | + | + |
| ***W40*** | + | + | + | + |
| ***LD*** | + | + | + | + |
| ***W10-LD*** | + | + | + | + |
| ***W20-LD*** | + | + | + | + |
| ***W40-LD*** | + | + | + | + |

<sup>1</sup>W10= categorical model with window of 10 SNPs, W20= categorical model with window of 20 SNPs, W40= categorical model with window of 40 SNPs, LD= continuous model with LD estimates, W10-LD= combined model with window of 10 SNPs and LD estimates, W20-LD= combined model with window of 20 SNPs and LD estimates, W40-LD= combined model with window of 40 SNPs and LD estimates.
<sup>2</sup>Sc0= simulation scenario 0, Sc1= simulation scenario 1, Sc2= simulation scenario 2, Sc3= simulation scenario 3.

### Hglm method and CodataGS

The estimation of the SNP effects was performed by fitting the model described by equations 1–3 that allows both continuous and categorical predictors for the SNP-specific variances, or a combination of continuous and categorical predictors. We tested a few examples of external information on the SNPs and these models were fitted using the **hglm** package in R (Rönnegård et al., 2010c). In the **hglm** package the linear predictor *α* + *βx_j_* is specified using the *X.rand.disp* option in the hglm function and the function computes fitted SNP effects (example of the command line to call the hglm function with the linear predictor option *X.rand.disp* can be found in the Supplementary File S1 line 156).

When the number of markers largely exceeds the number of individuals, the computational speed and memory requirements can be improved by fitting individual effects (i.e. EGBVs) in an equivalent model instead of SNP effects. This equivalent model, which uses the external information on each SNP in the same way as in the **hglm** package, was implemented in the R package **CodataGS** and is available in the Supplementary File S3. The theory is explained in the Supplementary File S4.

### Accuracy

The predictive ability of all models was evaluated as the correlation of the estimated genomic breeding values (EGBVs) and the true genomic breeding values (TGBVs) for the validation set (Generation 2). For each simulation setup, 100 replicates were generated. The convergence of the models varied from 71% to 100% and results are presented for those replicates where all models converged.

## Data Availability

Simulation of the data that support the findings is possible through the attached simulation code in File S1 and File S2 (Functions for the simulation) deposited at fig**share**. The simulation code and the methodology described previously are sufficient to reproduce the results of this study.

## Results

Table 1 contains different versions of the model tested. The fitted SNP effects obtained from **hglm** for one simulation replicate under scenario Sc0 with 10 QTL per chromosome are presented in Figure 1. The R code to reproduce Figure 1 is found in Supplementary File S1 (along with File S2). The results show how the fitted SNP effects may change between model specifications. For example, it can be observed that with increasing window size the estimated effects tend to be spread between more SNPs.

**Figure 1.**
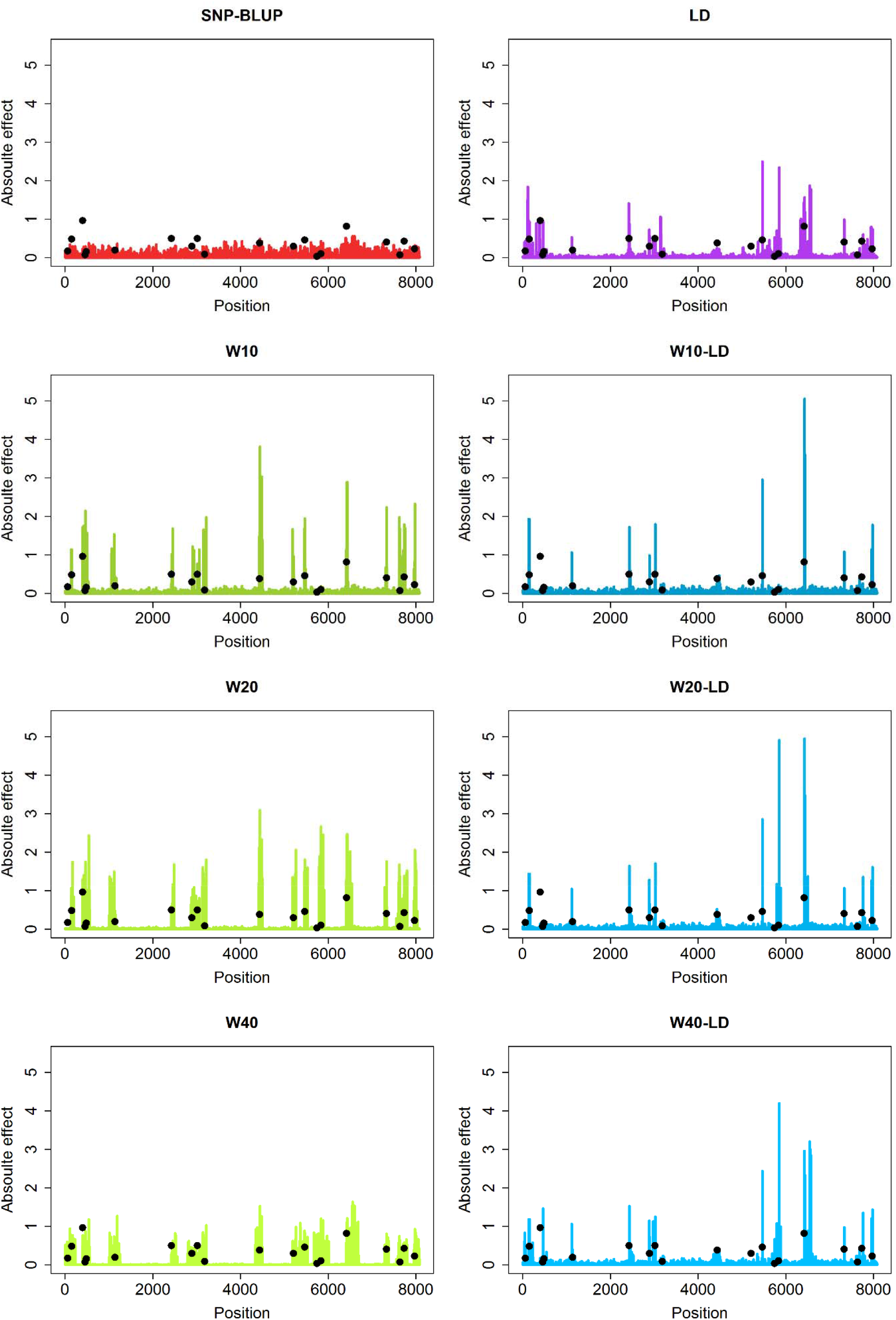
Fitted SNP effects under SNP-BLUP and 7 alternative models for one simulation replicate under simulation scenario Sc0 with 10 QTL per chromosome underlying the trait.

### Model performance

Table 2 shows the accuracies of the predicted EGBVs in the validation set (generation 2) for scenario 0 (Sc0) with 10 QTLs per chromosome underlying the trait. In general, the alternative models performed better than SNP-BLUP. The categorical models yielded higher accuracies compared to the SNP-BLUP model by 14.3% (0.670 ± 0.013), 11.9% (0.656 ± 0.012) and 8.4% (0.635 ± 0.012) for the models W10, W20 and W40, respectively. Nonetheless, it can be observed that the advantage of the categorical models over the SNP-BLUP decreased with increasing window sizes. Moreover, the continuous model (LD) resulted in higher accuracy than the SNP-BLUP or the categorical models with an increase of 22.4% (0.717 ± 0.011) in accuracy with respect to the SNP-BLUP. Similarly, the combination models performed 20.6% (W10-LD, 0.707 ± 0.013) 21.8% (W20-LD, 0.714 ± 0.013) and 23.2% (W40-LD, 0.722 ±0.013) better than the SNP-BLUP model. Contrary to the categorical models, the combination models maintained the gain in accuracy with increasing window size. Finally, the alternative models provided unbiased predictions while the SNP-BLUP showed upward bias (Table 2).

**Table 2.** Accuracy and bias of the predicted EGBVs in the validation set (generation 2) for the scenario 0 (Sc0) with 10 QTLs per chromosome underlying the trait.

| Models<sup>1</sup> | Accuracy (r) | Bias (b) |
| --- | --- | --- |
| **SNP-BLUP** | 0.586 <sup>(0.010)</sup> | 1.213 <sup>(0.089)</sup> |
| **W10** | 0.670 <sup>(0.013)</sup> | 1.003 <sup>(0.044)</sup> |
| **W20** | 0.656 <sup>(0.012)</sup> | 1.014 <sup>(0.048)</sup> |
| **W40** | 0.635 <sup>(0.012)</sup> | 1.030 <sup>(0.045)</sup> |
| **LD** | 0.717 <sup>(0.011)</sup> | 1.024 <sup>(0.041)</sup> |
| **W10-LD** | 0.707 <sup>(0.013)</sup> | 1.050 <sup>(0.053)</sup> |
| **W20-LD** | 0.714 <sup>(0.013)</sup> | 1.044 <sup>(0.050)</sup> |
| **W40-LD** | 0.722 <sup>(0.013)</sup> | 1.028 <sup>(0.042)</sup> |

<sup>1</sup>W10= categorical model with window of 10 SNPs, W20= categorical model with window of 20 SNPs, W40= categorical model with window of 40 SNPs, LD= continuous model with LD estimates, W10-LD= combined model with window of 10 SNPs and LD estimates, W20-LD= combined model with window of 20 SNPs and LD estimates, W40-LD= combined model with window of 40 SNPs and LD estimates.

### Effect of number of simulated QTLs

In order to investigate the performance of the alternative models for traits with different genetic architectures we simulated a trait controlled by an increasing number of QTLs with each having a decreasing effect. As an overview, the accuracies of the different models in Sc0 with 20 and 100 QTL per chromosome are visualized in Figure 2 together with the results from 10 QTL per chromosome. The advantage of the alternative models over the SNP-BLUP model decreased with increasing number of QTLs controlling the trait. When the number of QTLs underlying the trait is 20 QTLs per chromosome, the accuracies obtained were 9.6%, 7.5% and 3.8% better than the SNP-BLUP for the W10, W20 and W40 models, respectively. The continuous model resulted in a gain of 12.9% in accuracy while the combination models performed slightly better than all the alternative models yielding gains in accuracy of 14%, 14.2% and 13.5% for the W10-LD, W20-LD and W40-LD models, respectively. Finally, in the case of 100 QTLs per chromosome, all models performed roughly the same as SNP-BLUP, yielding accuracies between 0.583 ± 0.012 (W40) and 0.599 ± 0.011 (W10-LD).

**Figure 2.**
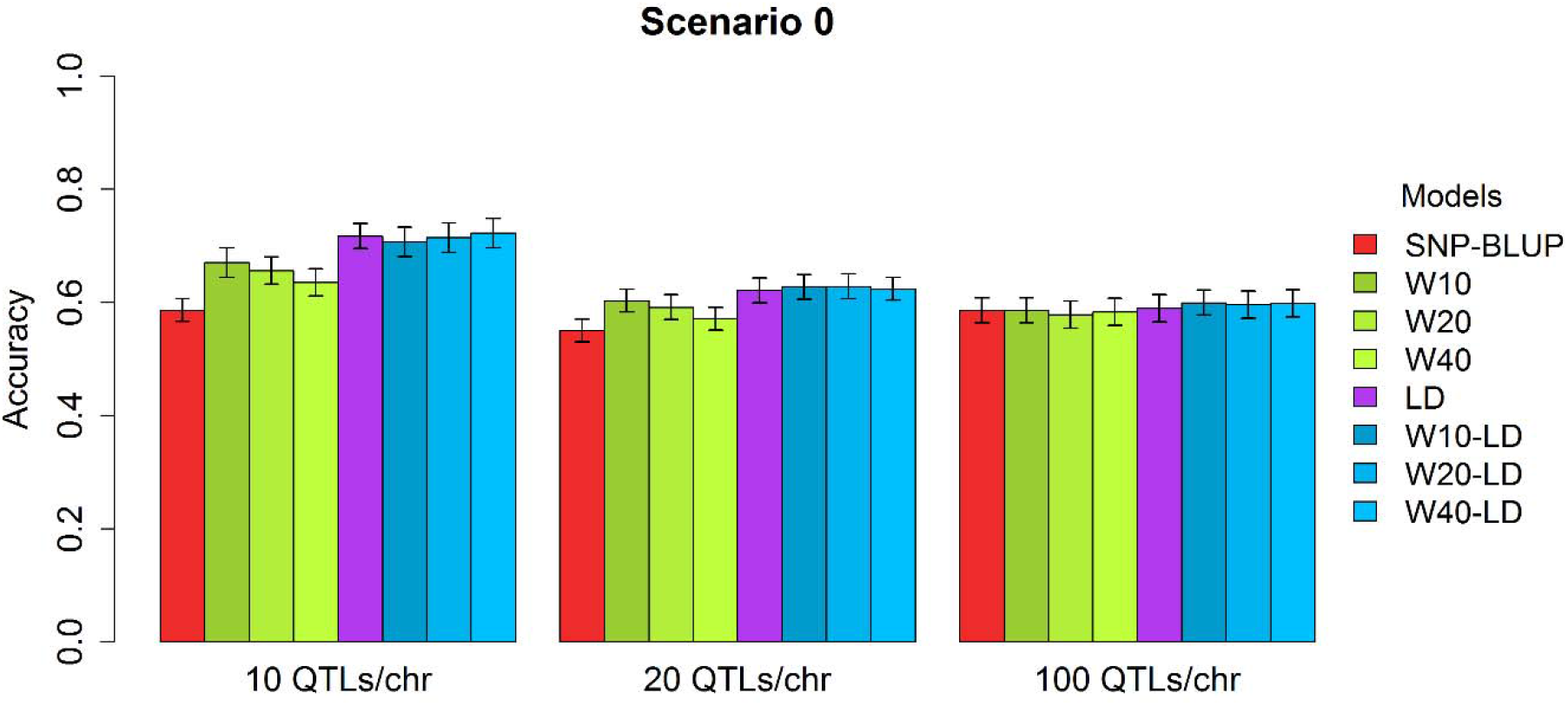
Accuracies obtained under different cases of genetic architecture of the trait for SNP-BLUP and the alternative models.

### Effect of variance of the QTL effects

The genetic architecture of a trait does not only depend on the number of QTLs that affect the trait. For example, mutations can affect protein coding regions or regulatory regions and these mutations can have a bigger or smaller effect on the trait. Therefore we can assume that their effects come from a mixture of distributions with varying variance over the genome. For this purpose we simulated several scenarios where the QTL effects were drawn from a mixture of distributions (see Sc1 – Sc3 in Materials and Methods). We compared the performance of all models under all scenarios of QTL effect variances and all cases of number of QTLs affecting the trait (Figure 3). In general the models performed similarly under Sc1, Sc2 and Sc3 as in Sc0. Small differences were observed in the case of 10 QTLs per chromosome where all models performed slightly better in Sc0 and Sc2 (QTL effects from a low variance distribution on chromosome 1 and from high variance distribution on chromosome 2) compared with the results from Sc1 and Sc3. Nonetheless, this minimum difference disappeared quickly with increasing number of QTLs per chromosomes. The external information included in the alternative models was related to the position of the QTLs on the genome and/or the relationship of the SNPs with the QTLs (LD), but no information about the distribution of the variance itself was included. Therefore, we fitted additional models that considered the way the QTLs were simulated (see linear predictor 5: Additional models Material and Methods, and Supplementary file Table S1). For Sc1 and Sc2 we extended the linear predictor (*α* + *βx*_*j*1_ + *γx*_*j*2_) to accommodate for two types of variances for the SNPs in windows that harboured a QTL assuming that we knew beforehand the distribution variance of the effect of that QTL and, as before, we tested 3 different window sizes (10, 20 and 40 SNPs per window). The results showed that these additional models performed similarly as the categorical models (W10, W20 and W40) under all cases of genetic architecture simulated. The only exception to these results was for the Sc2 with 100 QTLs per chromosome where additional models showed a small increase in accuracy compared to all other models (Supplementary files, Figure S1). For the Sc3 we used the distance of the SNP from the edge of the chromosome as external information, either as a continuous variable or within windows. Similarly as before, the additional models that included information on the simulated distribution variance of the QTLs did not perform better than the alternative models. The combined models (W10-Dis, W20-Dis and W40-Dis) performed the same as the categorical models while the continuous model (Dis) showed no benefit compared to the alternative models or the SNP-BLUP model under any simulation scenario of genetic architecture.

**Figure 3.**
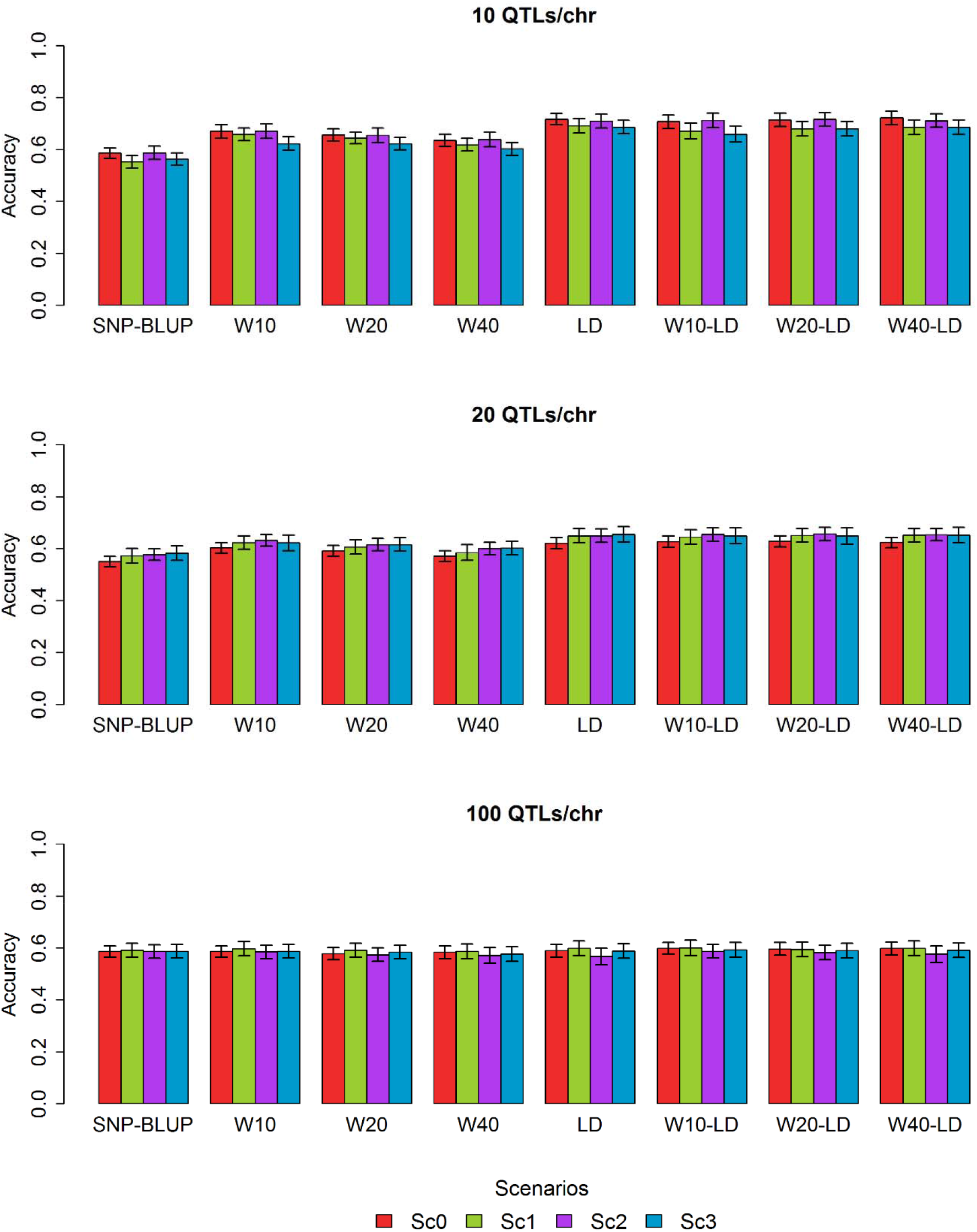
Accuracies obtained from SNP-BLUP model and alternative models under all simulated scenarios and genetic architectures.

### Computation time

When the number of markers exceeds the number of individuals, the computational speed and memory requirements can be an important drawback for the use of such models. A solution to this problem is to fit individual effects (i.e. EGBVs) in an equivalent model instead of SNP effects. In this study all evaluations were performed using the **hglm** R package that fits SNP effects. For a larger number of SNPs the computations would be unfeasible and an equivalent model which uses the external information on each SNP in the same way as in the **hglm** package was implemented in the R package **CodataGS** (Supplementary File S3). The theory is explained in the Supplementary File S4. Fitting individual effects instead of SNP effects resulted in largely improved run time of all models. For a training population of 200 individuals with 2,000 SNP markers, fitting SNP effects (**hglm**) required on average 9.35 seconds per iteration while fitting individual effects (**CodataGS**) required only 0.46 seconds per iteration (Figure 4). The improved speed and memory requirements of the equivalent model can be considerably beneficial since the usual size of the training sets is much larger than the one used here (thousands of individuals with tens of thousands of SNPs).

**Figure 4.**
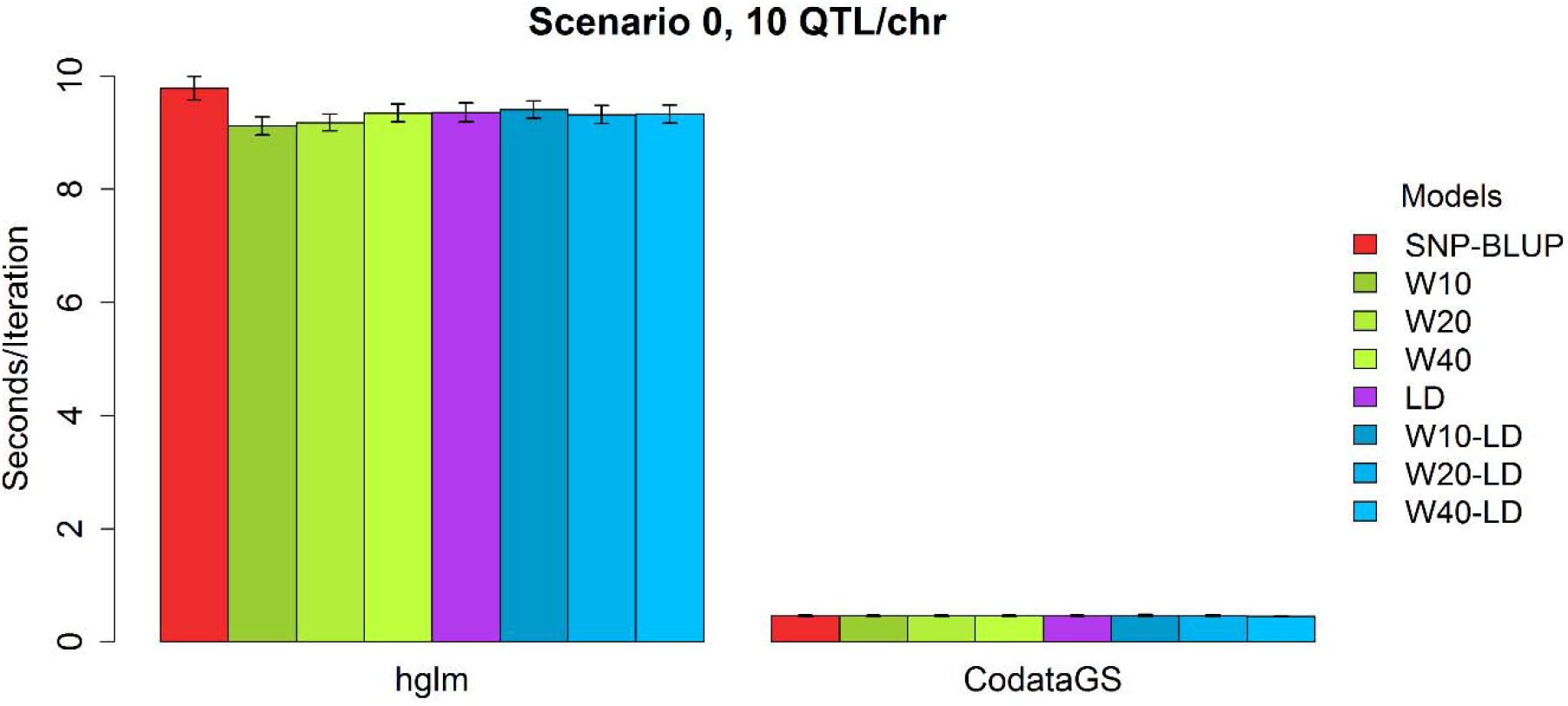
Time of execution (seconds per iteration) of SNP-BLUP and alternative models from **hglm** package and **CodataGS** package.

## Discussion

The knowledge on the genetic architecture of different traits, and SNP-specific biological information, is increasing rapidly and several authors have proposed methods for genomic selection that can make use of this available biological information to improve selection accuracy (Zhang et al., 2010; Zhang et al., 2014; Su et al., 2014). In this line, this study proposes a general model using a link function approach within the hierarchical generalized linear model framework (Lee et al., 2006) to include biological external information into the model.

All the results in the current study use the same general model (described by equations 1 – 3) for predicting genomic breeding values. The alternative models in Table 1, including SNP-BLUP, are fitted within this single framework and in the results the accuracies of the alternative models are compared. There are numerous Bayesian models not included within this framework that may be of interest to compare with. However, we use SNP-BLUP as a basic model to compare the results to and study the accuracies of models that make use of external information on the SNPs

A very attractive feature of the method proposed in this study is that it provides a flexible way to model the SNP variances using a linear predictor (eq. 3). Any type of existing knowledge on the SNP markers can be utilized and potentially increase the predictive ability of the model. In this study we investigated the performance of external information related to the position of the QTLs on the genome and the relationship of the SNP markers with the QTLs and we showed that the inclusion of such information can improve the predicting ability of genomic selection. From our results we identified two main factors that influence the performance of such models, the genetic architecture of the trait and the quality/accuracy of the external information.

### Genetic architecture of the trait

The performance of several alternative models in our study was better compared to the SNP-BLUP method when the trait was controlled by a small number of QTLs with medium-large effects. The advantage of these models was reduced with increasing number of QTLs with smaller effects. However, the alternative models did not result in lower accuracies compared to the SNP-BLUP model. The reason is that as the estimated effect of the external information on the SNP variances approaches zero the model reduces to a SNP-BLUP model. Furthermore, as the number of QTL that control the trait increases, the external information on SNPs becomes more similar among the SNPs. For example, for the categorical models, a QTL is located within most or all defined windows and as a result all SNPs get the same weight in the model. Moreover, most or all SNPs are in LD with a QTL at similar levels. Consequently, the alternative models turn into a SNP-BLUP model. These results are in agreement with the findings of Zhang et al., 2010. In their simulation study they investigated the performance of a BLUP model with weighted G matrix and showed that for traits controlled by high number of QTLs the traditional GBLUP and their method performed similarly. This effect has also been observed in studies on real data (Zhang et al., 2014). Analysing three dairy cattle traits (Milk Yield (MY), Fat percentage (FP) and Somatic Cell Count (SCC)) these authors found that traits controlled by a few QTLs with large effects (MY and FP) perform better under models with external information on the SNPs while the SCC trait, that is controlled by many QTLs evenly distributed along the genome, performed better under the standard GBLUP model.

In our simulation study we created different genetic architectures for the trait with respect not only to the number of the QTLs affecting the trait but also to the distribution of the QTL effects and their variances (see Material and Methods). Our results showed that this aspect did not affect the performance of the alternative models. Moreover, the additional models that included information on the variance distribution across the genome were not able to provide any benefit, contrary to methods that assume mixtures of distributions for the SNP markers like Bayesian methods (Erbe et al., 2012).

### External information

In this study we investigated the performance of models that include information on the location of the QTLs on the genome (categorical models) and thereby tried to mimic the external information available on the QTL databases and the different window sizes resemble the degree of uncertainty of a QTL region. Our results indicate that this type of external information has the potential to improve the accuracy of genomic selection and that the degree of improvement is inversely related to the degree of uncertainty on the QTL region. The usefulness of the QTL database information has been demonstrated by Zhang et al., 2014. In their study these authors searched for reported QTLs on the traits under consideration (Fat percentage, milk yield and somatic cell score for dairy cattle and several traits for rice) and after a quality control to avoid the possible false positive reports they included this information into a GBLUP model. For most of the examined traits an increase in accuracy was observed, especially for the traits that showed a characteristic genetic architecture. The discovery of new QTLs or the causative mutations is expected to increase in the future with the use of whole genome sequence and the development of new methods for analysis and as a consequence the information available will become more accurate.

The external information that proved to be more valuable in this study was the LD estimates between the SNPs and the QTLs. In the standard GBLUP method, markers in linkage equilibrium (LE) to the causative QTL tend to capture effects due to family relationship, whereas mainly markers in LD capture the QTL effects themselves (Habier et al. 2007, de los Campos et al. 2015). In the BayesB model (Meuwissen et al. 2001), the prior for the SNP variances is a mixture of two distributions that tends to group markers into two classes: those in LD and those in LE with the QTL. By modelling the two classes of markers better predictions for unrelated individuals can be obtained. In other studies, LD information has been incorporated in a model for the marker variances, which smooths the effects between markers in close LD (e.g. the Bayesian antedependence model by Yang and Tempelman 2012, and the double hierarchical generalized linear model by Rönnegård and Lee 2010b), and thereby captures the QTL effects rather than family information. These models give better predictions than GBLUP when individuals are unrelated and the total number of QTL is small. This is in line with our findings where the models including LD between markers and QTL resulted in improved prediction accuracies, especially when the number of simulated QTL was small. Finally, the results obtained from the combined models indicate that information on the real relationship between markers and QTLs can compensate for the loss of information due to the uncertainty of the QTL report.

## Conclusions

In this study we investigated the potential benefit of external information on improving the accuracy of genomic selection. In conclusion, using external information to model SNP-specific variances can provide gains in accuracy compared to the traditional SNP-BLUP. Nonetheless, the level of gain depends on the genetic architecture of the trait of interest and the quality of the external information on the SNP markers. The usefulness of these type of models is expected to increase with time as more accurate information on the SNPs becomes available. Finally, further studies are required to confirm the advantage of this approach on real data.

## Acknowledgements

This project was supported by the Mistra Biotech project, a research program financed by Mistra – the Swedish foundation for strategic environmental research, and the Swedish University of Agricultural Sciences, SLU. M. Selle acknowledges the financial support given by the Research Council of Norway, grant number 250362.

## References

Abdollahi-Arpanahi, R., G. Morota, B. D. Valente, A. Kranis, G. J. M. Rosa, et al., 2016 Differential contribution of genomic regions to marked genetic variation and prediction of quantitative traits in broiler chickens. Genet Sel Evol : GSE 48(1), 10.

Bush, W. S., and J. H. Moore, 2012 Chapter 11: Genome-Wide Association Studies. PLoS Comput Biol. 8(12):e1002822.

Caspi, R., T. Altman, K. Dreher, C. A. Fulcher, P. Subhraveti, et al., 2012 The MetaCyc database of metabolic pathways and enzymes and the BioCyc collection of pathway/genome databases. Nucleic Acids Res. 40(June), D742–D753.

Croft, D., G. O’Kelly, G. Wu, R. Haw, M. Gillespie, et al., 2011 Reactome: A Database of Reactions, Pathways and Biological Processes. Nucleic Acids Res. 39 (SUPPL. 1).

Daetwyler, H. D., R. Pong-Wong, B. Villanueva, and J. A. Woolliams, 2010 The impact of genetic architecture on genome-wide evaluation methods. Genetics, 185(3):1021–1031.

Daetwyler, H. D., M. P. L. Calus, R. Pong-Wong, G. de los Campos, and J. M. Hickey, 2013 Genomic Prediction in Animals and Plants: Simulation of Data, Validation, Reporting, and Benchmarking. Genetics 193 (2): 347 LP–365.

Daetwyler H. D., A. Capitan, H. Pausch, P. Stothard, R. van Binsbergen, et al., 2014 Whole-genome sequencing of 234 bulls facilitates mapping of monogenic and complex traits in cattle. Nat. Genet. 46(8):858–865.

de Los Campos, G., H. Naya, D. Gianola, J. Crossa, A. Legarra, et al., 2009 Predicting quantitative traits with regression models for dense molecular markers and pedigree. Genetics 182(1), 375–385.

de los Campos, G., D. Sorensen, and D. Gianola, 2015 Genomic heritability: What is it? PLOS Genetics, 11(5): e1005048.

Do, D. N., L. L. G. Janss, J. Jensen, and H. N. Kadarmideen, 2015 SNP Annotation-Based Whole Genomic Prediction and Selection : An Application to Feed Efficiency and Its Component Traits in Pigs. J Anim Sci. 93: 2056–63.

Erbe, M., B. J. Hayes, L. K. Matukumalli, S. Goswami, P. J. Bowman, et al., 2012 Improving accuracy of genomic predictions within and between dairy cattle breeds with imputed high-density single nucleotide polymorphism panels. J Dairy Sci. 95(7), 4114–4129.

Gao, N., J. Li, J. He, G. Xiao, Y. Luo, et al., 2015 Improving accuracy of genomic prediction by genetic architecture based priors in a Bayesian model. BMC Genet. 16:120.

Gianola, D., 2013 Priors in whole-genome regression: The Bayesian alphabet returns. Genetics 194(3), 573–596.

González-Recio, O., D. Gianola, G. J. M. Rosa, K. A. Weigel, and A. Kranis, 2009 Genome-Assisted Prediction of a Quantitative Trait Measured in Parents and Progeny: Application to Food Conversion Rate in Chickens. Genet Sel Evol : GSE 41 (1): 3.

Gunderson, K. L., F. J. Steemers, G. Lee, L. G. Mendoza and M. S. Chee, 2005 A genome-wide scalable SNP genotyping assay using microarray technology. Nat. Genet. 37:549–554.

Habier, D., R. L. Fernando, and J. C. M. Dekkers, 2007 The impact of genetic relationship information on genome-assisted breeding values. Genetics 177: 2389–2397.

Habier, D., R. L. Fernando, K. Kizilkaya, and D. J. Garrick, 2011 Extension of the bayesian alphabet for genomic selection. BMC Bioinformatics, 12(1), 186.

Hayes, B. J., P. J. Bowman, A. J. Chamberlain, and M. E. Goddard, 2009 Invited review: Genomic selection in dairy cattle: progress and challenges. J. Dairy Sci. 92(2):433–43.

Hayes, B. J., J. Pryce, A. J. Chamberlain, P. J. Bowman and M. E. Goddard, 2010 Genetic Architecture of Complex Traits and Accuracy of Genomic Prediction: Coat Colour, Milk-Fat Percentage, and Type in Holstein Cattle as Contrasting Model Traits. PLOS Genetics 6(9): e1001139.

Hecker, M., S. Lambeck, S. Toepfer, E. van Someren, and R. Guthke, 2009 Gene regulatory network inference: Data integration in dynamic models-a review. Biosystems 96:86–103.

Hidalgo, A. M., J. W. M. Bastiaansen, M. S. Lopes, B. Harlizius, M. A. M. Groenen, et al., 2015 Accuracy of predicted genomic breeding values in purebred and crossbred pigs. G3-GENES GENOM GENET 5: 1575–1583.

Hu, Z. L., C. A. Park, X. L. Wu, and J. M. Reecy, 2013 Animal QTLdb: An improved database tool for livestock animal QTL/association data dissemination in the post-genome era. Nucleic Acids Res. 41(D1), D871–D879.

Kadarmideen H., 2014 Genomics to systems biology in animal and veterinary sciences: Progress, lessons and opportunities. Livest. Sci. 166: 232–248.

Kanehisa, M., M. Araki, S. Goto, M. Hattori, M. Hirakawa, et al., 2008 KEGG for Linking Genomes to Life and the Environment. Nucleic Acids Res. 36 (SUPPL. 1).

Koufariotis, L., Y. P. Chen, S. Bolormaa, B. J. Hayes, A. J. Schork, et al., 2014 Regulatory and Coding Genome Regions Are Enriched for Trait Associated Variants in Dairy and Beef Cattle. BMC Genomics 15 (1). BioMed Central: 436.

Lee, T. I., N. J. Rinaldi, F. Robert, D. T. Odom, Z. Bar-Joseph, et al., 2002 Transcriptional regulatory networks in Saccharomyces cerevisiae. Science 298: 799–804

Lee, Y., and J. A. Nelder, 1996 Hierarchical Generalized Linear Models. J. R. Statist. Soc. B 58 (4).

Lee, Y., J. A. Nelder, and Y. Pawitan, 2006 Generalized Linear Models with Random Effects: Unified Analysis via H-likelihood. Chapman & Hall/CRC, Boca Raton. 80pp.

Legarra, A., C. Robert-Granié, E. Manfredi, and J. M. Elsen, 2008 Performance of Genomic Selection in Mice. Genetics 180 (1):611–18.

Luan, T., J. A. Woolliams, S. Lien, M. Kent, M. Svendsen, and T. H. E. Meuwissen, 2009 The Accuracy of Genomic Selection in Norwegian Red Cattle Assessed by Cross-Validation. Genetics 183 (3):1119–26.

MacLeod, I. M., P. J. Bowman, C. J. Vander Jagt, M. Haile-Mariam, K. E. Kemper, et al., 2016 Exploiting biological priors and sequence variants enhances QTL discovery and genomic prediction of complex traits. BMC Genomics, 17(1), 144.

Meuwissen, T. H. E., B. J. Hayes, and M. E. Goddard, 2001 Prediction of total genetic value using genome-wide dense marker maps. Genetics, 157(4), 1819–1829.

Morota, G., R. Abdollahi-Arpanahi, A. Kranis, and D. Gianola, 2014 Genome-Enabled Prediction of Quantitative Traits in Chickens Using Genomic Annotation. BMC Genomics 15 (1). BioMed Central: 109.

Muir, W. M., 2007 Comparison of Genomic and Traditional BLUP-Estimated Breeding Value Accuracy and Selection Response under Alternative Trait and Genomic Parameters. J. Anim. Breed. Genet. 124 (6): 342–55.

Ostersen, T., O. F. Christensen, M. Henryon, B. Nielsen, G. Su, and P. Madsen, 2011 Deregressed EBV as the Response Variable Yield More Reliable Genomic Predictions than Traditional EBV in Pure-Bred Pigs. Genet Sel Evol : GSE 43 (1): 38.

Rönnegård, L., M. Felleki, F. Fikse, H. A. Mulder, and E. Strandberg, 2010a. Genetic heterogeneity of residual variance – estimation of variance components using double hierarchical generalized linear models. Genet Sel Evol : GSE 42:8.

Rönnegård, L., and Y. Lee, 2010b Hierarchical generalized linear models have a great potential in genetics and animal breeding. In proceedings: World Congress on Genetics Applied to Livestock Production, Leipzig, Germany.

Rönnegård, L., X. Shen, and M. Alam, 2010c hglm: A Package for Fitting Hierarchical Generalized Linear Models. The R J., 2(2), 20–28.

Shalgi, R., D. Lieber, M. Oren, and Y. Pilpel, 2007 Global and local architecture of the mammalian microRNA-transcription factor regulatory network. PLOS Comput. Biol. 3:e131.

Snelling, W. M., R. A. Cushman, J. W. Keele, C. Maltecca, M. G. Thomas, et al., 2013 BREEDING AND GENETICS SYMPOSIUM: Networks and pathways to guide genomic selection. J. Anim. Sci. 91:537–552.

Sonesson, A. K., and T. H. E. Meuwissen, 2009 Testing Strategies for Genomic Selection in Aquaculture Breeding Programs. Genet Sel Evol: GSE 41:37.

Su, G., O. F. Christensen, L. Janss and M. S. Lund, 2014 Comparison of genomic predictions using genomic relationship matrices built with different weighting factors to account for locus/specific variances. J. Dairy Sci. 97:6547–6559

Tusell, L., H. Gilbert, J. Riquet, M. J. Mercat, A. Legarra, et al., 2016. Pedigree and genomic evaluation of pigs using a terminal-cross model. Genet Sel Evol: GSE 48:32

Usai, M. G., M. E. Goddard, and B. J. Hayes, 2009 LASSO with cross-validation for genomic selection. Genetics Research, 91(6), 427–436.

VanRaden, P. M., C. P. Van Tassell, G. R. Wiggans, T. S. Sonstegard, R. D. Schnabel, et al., 2009 Invited Review: Reliability of Genomic Predictions for North American Holstein Bulls. J Dairy Sci. 92 (1):16–24.

Wolc, A., H. H. Zhao, J. Arango, P. Settar, J. E. Fulton, et al., 2015 Response and inbreeding from a genomic selection experiment in layer chickens. Genet Sel Evol: GSE 47(1):59.

Yang, J., T. A. Manolio, L. R. Pasquale, E. Boerwinkle, N. Caporaso, et al., 2011 Genome Partitioning of Genetic Variation for Complex Traits Using Common SNPs. Nature Genetics 43 (6): 519–25.

Yang, W. and R. J. Tempelman, 2012 A Bayesian antedependence model for whole genome prediction. Genetics: 1491–1501.

Zhang, Z., J. F. Liu, X. D. Ding, P. Bijma, D. J. de Koning, et al., 2010 Best linear unbiased prediction of genomic breeding values using trait-specific marker-derived relationship matrix. PLoS ONE 5: e12648

Zhang Z, U. Ober, M. Erbe, H. Zhang, N. Gao, et al., 2014 Improving the accuracy of Whole Genome Prediction for Complex Traits using the results of Genome Wide Association Studies. PLoS ONE 9(3): e93017.

