## Supplementary File S1 for "Genomic prediction including SNP-specific variance predictors"

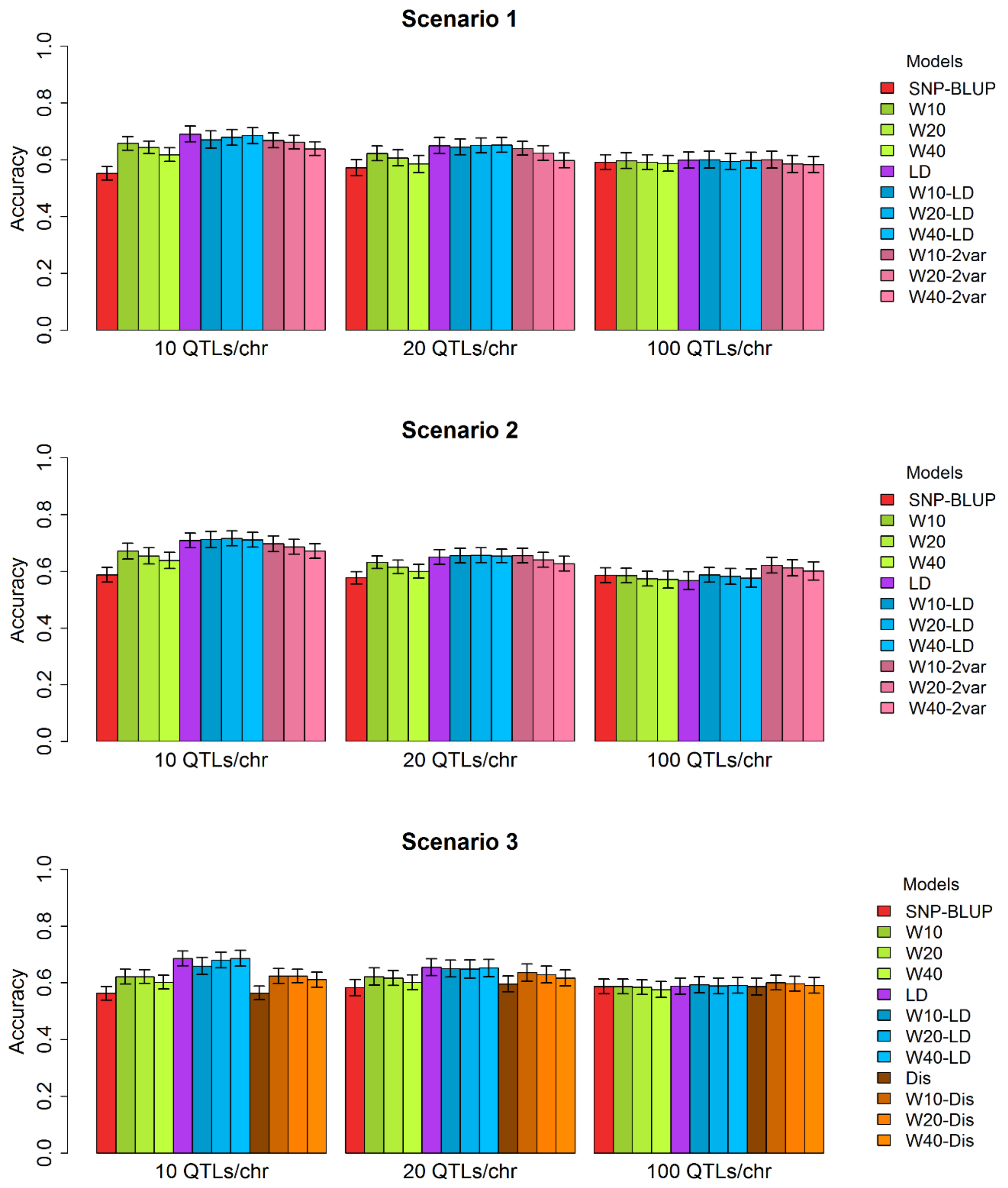


**Figure S1.** Accuracies (standard errors) obtained from all models tested (SNP-BLUP, alternative and additional models) under scenarios of simulated genetic architecture 1, 2 and 3.
