## Supplementary File S4 for "Genomic prediction including SNP-specific variance predictors"

### Supplementary material: Equivalent models

To derive a computationally efficient implementation of a regression algorithm exploiting codata for SNP markers for $p \gg n$, we present equivalent models to the SNP-BLUP and show how the estimates of the effects, and their associated prediction error variances, can be transformed between these. Prediction error variances are important to compute since they are the basis for calculations of standard errors and effective number of parameters $p_{eff}$ [2, 3]. Most of theory below is found in [10] but here the focus is on fitting a regression model for SNP-specific variances using codata on the SNP markers, and some minor typos of [10] have been corrected below.

Three different, but equivalent, specifications of the random effects will be used and will be referred to as the *SNP Model*, the *animal model* and the *Cholesky Model*. The SNP effects may have variances with different weights given by the diagonal matrix $\boldsymbol{D}$, and the residuals $\boldsymbol{e}$ are assumed iid normal with variance $\sigma_e^2$:

**SNP Model**

$$\begin{aligned} \mathbf{y} &= \mathbf{X}\boldsymbol{\beta} + \mathbf{Z}\mathbf{b} + \mathbf{e} \\ \mathbf{b} &\sim N(0, \sigma_b^2 \mathbf{D}) \end{aligned} \quad (1)$$

**Animal model**

$$\begin{aligned} \mathbf{y} &= \mathbf{X}\boldsymbol{\beta} + \mathbf{a} + \mathbf{e} \\ \mathbf{a} &\sim N(0, \sigma_b^2 \mathbf{G}) \\ \mathbf{G} &= \mathbf{Z}\mathbf{D}\mathbf{Z}' \end{aligned} \quad (2)$$

**Cholesky Model**

$$\begin{aligned} \mathbf{y} &= \mathbf{X}\boldsymbol{\beta} + \mathbf{L}\mathbf{v} + \mathbf{e} \\ \mathbf{v} &\sim N(0, \sigma_b^2 \mathbf{I}_n) \\ \mathbf{L}\mathbf{L}' &= \mathbf{G} \end{aligned} \quad (3)$$

The use of equivalent LMMs in the research field of animal breeding and quantitative genetics is well established. The contribution here is to present how LMM theory can be used for heteroscedastic variance components, to show how the prediction error variances can be transformed between models, to implement the theory in a computationally efficient R package **CodataGS**, and to apply it to a model including external codata on the SNP genotypes presented further below.

Different mixed model equations for the equivalent models

For LMM Henderson's mixed model equations (MME) are used to estimate both the fixed and random effects for given variance components. They can also be used iteratively to estimate variance components as implemented in the R package **hglm** [9]. Although the models above are equivalent, the MME are different.

**SNP model**

For the SNP Model we have the MME

$$\begin{pmatrix} \mathbf{X}'\mathbf{X} & \mathbf{X}'\mathbf{Z} \\ \mathbf{Z}'\mathbf{X} & \mathbf{Z}'\mathbf{Z} + \frac{\sigma_e^2}{\sigma_b^2}\mathbf{D}^{-1} \end{pmatrix} \begin{pmatrix} \boldsymbol{\beta} \\ \mathbf{b} \end{pmatrix} = \begin{pmatrix} \mathbf{X}'\mathbf{y} \\ \mathbf{Z}'\mathbf{y} \end{pmatrix} \quad (4)$$

These MMEs are of size $(k + p) \times (k + p)$, where $k$ is the number of columns in $\boldsymbol{X}$. Hence, the size of the equations are very large for high-dimensional data.

**Animal model**

Let the random effects $\boldsymbol{a}$ be individual effects for each observation and $\boldsymbol{G} = \boldsymbol{ZDZ}'$ the correlation matrix between these. Then $\boldsymbol{G}$ is relatively small ($n \times n$) and the MME are

$$\begin{pmatrix} \mathbf{X}'\mathbf{X} & \mathbf{X}' \\ \mathbf{X} & \mathbf{I} + \frac{\sigma_e^2}{\sigma_b^2}\mathbf{G}^{-1} \end{pmatrix} \begin{pmatrix} \boldsymbol{\beta} \\ \mathbf{a} \end{pmatrix} = \begin{pmatrix} \mathbf{X}'\mathbf{y} \\ \mathbf{y} \end{pmatrix} \quad (5)$$

of size $(k + n) \times (k + n)$. Hence, the size of these MME is much smaller than eq. 4 for

$p \gg n$.

**Cholesky model**

In a third equivalent model we define $\mathbf{LL}' = \mathbf{G}$ (where $\mathbf{L}$ has size $n \times n$ ) and the random effects $\boldsymbol{v}$ are individual independent random effects. The MME are

$$\begin{pmatrix} \mathbf{X}'\mathbf{X} & \mathbf{X}'\mathbf{L} \\ \mathbf{L}'\mathbf{X} & \mathbf{L}'\mathbf{L} + \frac{\sigma_e^2}{\sigma_b^2}\mathbf{I}_n \end{pmatrix} \begin{pmatrix} \boldsymbol{\beta} \\ \mathbf{v} \end{pmatrix} = \begin{pmatrix} \mathbf{X}'\mathbf{y} \\ \mathbf{L}'\mathbf{y} \end{pmatrix} \tag{6}$$

of size $(k+n) \times (k+n)$.

Transformation of effects between equivalent models

For $p \gg n$, the size of the MME in models (5) and (6) are much smaller than in model (4).

The random effects can be transformed between these equivalent models [6, 7] so that the estimated SNP effects $\widehat{\boldsymbol{b}}$ can easily be calculated from the individual effects $\widehat{\boldsymbol{a}}$ in model (5)

$$\widehat{\boldsymbol{b}} = \mathbf{D}\mathbf{Z}'\mathbf{G}^{-1}\hat{\mathbf{a}}. \tag{7}$$

Furthermore, we have $\widehat{\boldsymbol{a}} = \mathbf{L}\hat{\mathbf{v}}$ so that

$$\widehat{\boldsymbol{b}} = \mathbf{D}\mathbf{Z}'\mathbf{G}^{-1}\mathbf{L}\hat{\mathbf{v}}. \tag{8}$$

The matrix $\mathbf{Z}$ is moderately large ($n \times p$) but the transformation is a simple cross-product.

Hence, the calculations can be made in parts without reading all of $\mathbf{Z}$ into memory. They can also easily be parallelized if necessary.

Transformation of prediction error variances between equivalent models

Not only the estimates, but also the prediction error variances (*i.e.* the diagonal elements of $Var(\mathbf{v} - \hat{\mathbf{v}}|\mathbf{v})$), are important to compute to allow for model checking and inference.

In the Cholesky model (eq. 6), let $\mathbf{C_v}$ be

$$\mathbf{C_v} = \frac{1}{\sigma_e^2}\begin{pmatrix} \mathbf{X}'\mathbf{X} & \mathbf{X}'\mathbf{L} \\ \mathbf{L}'\mathbf{X} & \mathbf{L}'\mathbf{L} + \frac{\sigma_e^2}{\sigma_b^2}\mathbf{I}_n \end{pmatrix}. \tag{9}$$

Decompose the inverse of $\mathbf{C_v}$ as

$$\mathbf{C_v}^{-1} = \begin{pmatrix} \mathbf{C_v}^{11} & \mathbf{C_v}^{12} \\ \mathbf{C_v}^{21} & \mathbf{C_v}^{22} \end{pmatrix}. \tag{10}$$

Then the prediction covariance matrix is $Var(\mathbf{v}-\hat{\mathbf{v}}|\mathbf{v}) = \sigma_b^2\mathbf{I}_n - \mathbf{C}_{\mathbf{v}}^{22}$, see [1].

Denote the $j$:th diagonal element of $Var(\mathbf{b}-\hat{\mathbf{b}}|\mathbf{b})$ as $V_{\hat{b}_j}$. Then these elements can be calculated separately as

$$V_{\hat{b}_j} = \sigma_b^2 - \mathbf{M}_j(\sigma_b^2\mathbf{I}_n - \mathbf{C}_{\mathbf{v}}^{22})\mathbf{M}_j'\mathbf{D}^{-1} \tag{11}$$

where $\mathbf{M}_j$ is the $j$:th row of the transformation matrix $\mathbf{M} = \mathbf{DZ}'\mathbf{G}^{-1}\mathbf{L}$.

These computations are derived as follows. The prediction covariance matrix is

$$Var(\mathbf{b}-\hat{\mathbf{b}}|\mathbf{b}) = \sigma_b^2\mathbf{D} - \mathbf{C}_{\mathbf{b}}^{22} \tag{12}$$

$\mathbf{M}$ is the matrix transforming effects $\mathbf{v}$ to $\boldsymbol{b}$ in eq. (8), then

$$Var(\mathbf{b}-\hat{\mathbf{b}}|\mathbf{b}) = \mathbf{M}Var(\mathbf{v}-\hat{\mathbf{v}}|\mathbf{v})\mathbf{M}' \tag{13}$$

Combining these two equations, we get

$$\sigma_b^2\mathbf{D} - \mathbf{C}_{\mathbf{b}}^{22} = \mathbf{M}Var(\mathbf{v}-\hat{\mathbf{v}}|\mathbf{v})\mathbf{M}' \tag{14}$$

*i.e.*

$$\sigma_b^2\mathbf{D} - \mathbf{C}_{\mathbf{b}}^{22} = \mathbf{M}(\sigma_b^2\mathbf{I}_n - \mathbf{C}_{\mathbf{v}}^{22})\mathbf{M}' \tag{15}$$

So

$$\mathbf{C}_{\mathbf{b}}^{22} = \sigma_b^2\mathbf{D} - \mathbf{M}(\sigma_b^2\mathbf{I}_n - \mathbf{C}_{\mathbf{v}}^{22})\mathbf{M}' \tag{16}$$

<u>A simple relationship between hat values for random effects and the prediction error variance</u>

Henderson's MME are equivalent to the solving an augmented model using weighted least squares [4, 8]. The augmented design matrix for the SNP model is

$$\mathbf{T} = \begin{pmatrix} \mathbf{X} & \mathbf{Z} \\ \mathbf{0} & \mathbf{I} \end{pmatrix} \tag{17}$$

and the weight matrix is

$$\mathbf{W} = \begin{pmatrix} \frac{1}{\sigma_e^2}\mathbf{I} & \mathbf{0} \\ \mathbf{0} & \frac{1}{\sigma_b^2}\mathbf{D}^{-1} \end{pmatrix} \tag{18}$$

and the *hat matrix* for this augmented model is $\mathbf{H} = \mathbf{T}(\mathbf{T}'\mathbf{W}\mathbf{T})^{-1}\mathbf{T}'\mathbf{W}$.

Let $\mathbf{H}_{bb}$ be the lower right $p \times p$ part of $\mathbf{H}$ and $\mathbf{H}_{vv}$ the corresponding submatrix for the Cholesky model. Then the relationship between these submatrices and the prediction error variances are:

$$\mathbf{H}_{vv} = \mathbf{C_v}^{22}\frac{1}{\sigma_b^2} \tag{19}$$

and

$$\mathbf{H}_{bb} = \mathbf{C_b}^{22}\mathbf{D}^{-1}\frac{1}{\sigma_b^2} \tag{20}$$

Consequently, the hat values for the SNP model (i.e. the *j*:th diagonal element in $\mathbf{H}_{bb}$, $\mathrm{h}_{jj}$) can be computed efficiently from the prediction error variance of the Cholesky model.

$$h_{jj} = 1 - \mathbf{M}_j(\mathbf{I}_n - \mathbf{C}_{\mathbf{v}}^{22}/\hat{\sigma}_b^2)\mathbf{M}_j'\mathbf{D}_{jj}^{-1} \tag{21}$$

where $\mathbf{M}_j$ is the *j*:th row of the transformation matrix $\mathbf{M}$ and $\mathbf{D}_{jj}^{-1}$ is the j:th diagonal element of $\mathbf{D}^{-1}$.

Let $h_{bb}$ be the vector of the diagonal elements in $\mathbf{H}_{bb}$. In the R package **CodataGS**, $h_{bb}$ is efficiently computed as $h_{bb} = \mathbf{1} - \mathbf{c}_{sum}\mathbf{D}^{-1}$, where $\mathbf{c}_{sum}$ is the sum of columns of $(\mathbf{PM}') \circ (\mathbf{PM}')$, $\circ$ is the direct (elementwise) Hadamard product, and $\mathbf{P}$ is a square root matrix of

$\mathbf{I}_n - \mathbf{C}_{\mathbf{v}}^{22}/\hat{\sigma}_b^2$ computed using Singular Value Decomposition.

<u>Fitting SNP variances using codata with a generalized linear model</u>

Let $\boldsymbol{X}_{codata}$ be a design matrix with $p$ rows including covariates for the SNP markers. Then marker specific variances can be computed using a generalized linear model with a Gamma distribution and a log link function with response values $\frac{b^2}{1-h_{bb}}$ and weights $\frac{1-h_{bb}}{2}$ [4, 8], and linear predictor $\boldsymbol{X}_{codata}\boldsymbol{\beta}_{codata}$. An iterative procedure is applied where the diagonal elements in $\boldsymbol{X}_{codata}$ are updated using the fitted values: $\exp(\boldsymbol{X}_{codata}\widehat{\boldsymbol{\beta}}_{codata})$. This is the iterative reweighted least squares procedure for hierarchical generalized linear models derived in [3] and presented in [4] and [8].

#### References


[1] C. R. Henderson. *Applications of linear models in animal breeding*. University of Guelph, Guelph Ontario, 1984.

[2] D. J. Spiegelhalter, N. G. Best, B. P. Carlin, and A. Van Der Linde. Bayesian measures of model complexity and fit. *Journal of the Royal Statistical Society B*, 64(4):583-639.

[3] Y. Lee and J. A. Nelder. Hierarchical generalized linear models (with discussion). *Journal of the Royal Statistical Society B*, 58:619-678, 1996.

[4] Y. Lee, J. A. Nelder, and Y. Pawitan. *Generalized linear models with random effects - unified analysis via h-likelihood*. Chapman & Hall/CRC, 2006.

[5] Y. Lee, L. Rönnegård, and M. Noh. *Data Analysis Using Hierarchical Generalized Linear Models with R*. Chapman and Hall/CRC, 2017.

[6] M. Lynch and B. Walsh. *Genetics and analysis of Quantitative Traits*. Sinauer Associates, Inc., 1998.

[7] Y. Nagamine. Transformation of QTL genotypic effects to allelic effects. *Genetics Selection Evolution*, 37:579-584, 2005.

[8] L. Rönnegård, M. Felleki, F. Fikse, H. A. Mulder, and E. Strandberg. Genetic heterogeneity of residual variance - estimation of variance components using double hierarchical generalized linear models. *Genetics Selection Evolution*, 42:8, 2010.

[9] L. Rönnegård, X. Shen, and M. Alam. hglm: A package for fitting hierarchical generalized linear models. *The R Journal,* 2(2):20-28, 2010.

[10] X. Shen, M. Alam, F. Fikse, and L. Rönnegård. A novel generalized ridge regression method for quantitative genetics. *Genetics*, 193:1255-1268, 2013.
